# The systematic assessment of completeness of public metadata accompanying omics studies in the Gene Expression Omnibus

**DOI:** 10.1101/2021.11.22.469640

**Authors:** Yu-Ning Huang, Pooja Vinod Jaiswal, Anushka Rajesh, Anushka Yadav, Dottie Yu, Fangyun Liu, Grace Scheg, Emma Shih, Grigore Boldirev, Irina Nakashidze, Aditya Sarkar, Jay Himanshu Mehta, Ke Wang, Khooshbu Kantibhai Patel, Mustafa Ali Baig Mirza, Kunali Chetan Hapani, Qiushi Peng, Ram Ayyala, Ruiwei Guo, Shaunak Kapur, Tejasvene Ramesh, Dumitru Ciorbă, Viorel Munteanu, Viorel Bostan, Mihai Dimian, Malak S. Abedalthagafi, Serghei Mangul

## Abstract

Recent advances in high-throughput sequencing technologies have made it possible to collect and share a massive amount of omics data, along with its associated metadata. Enhancing metadata availability is critical to ensure data reusability and reproducibility and to facilitate novel biomedical discoveries through effective data reuse. Yet, incomplete metadata accompanying public omics data may hinder reproducibility and reusability by reducing sample interpretability and limiting secondary analyses. In this study, we performed a comprehensive assessment of metadata completeness shared in both scientific publications and/or public repositories by analyzing over 253 studies encompassing over 164 thousands samples, including both human and non-human mammalian studies. We observed that studies often omit over a quarter of important phenotypes, with an average of only 74.8% of them shared either in the text of publication or the corresponding repository. Notably, public repositories alone contained 62% of the metadata, surpassing the textual content of publications by 3.5%. Only 11.5% of studies completely shared all phenotypes, while 37.9% shared less than 40% of the phenotypes. Studies involving non-human samples were more likely to share metadata than studies involving human samples. We observed similar results on the extended dataset spanning 2.1 million samples across over 61,000 studies from the Gene Expression Omnibus repository. The limited availability of metadata reported in our study emphasizes the necessity for improved metadata sharing practices and standardized reporting. Finally, we discuss the numerous benefits of improving the availability and quality of metadata to the scientific community and beyond, supporting data-driven decision-making and policy development in the field of biomedical research. This work provides a scalable framework for evaluating metadata availability and may help guide future policy and infrastructure development.

## Introduction

Advancements in the high throughput sequencing technologies over the last decade have made omics data readily available to the public, enabling researchers to access a vast array of data across various diseases and phenotypes from the textual content of publications or various public repositories^1^. The vast amount of omics data and its accompanying metadata enable unprecedented exploration of biological systems through the re-analysis of public omics data, which is also known as secondary analysis^2,3^. Secondary analysis is capable of driving transformative breakthroughs in biomedical research, fostering collaboration, accelerating scientific progress, and deepening the understanding of human biology and diseases. By leveraging this readily available wealth of omics information, researchers can unravel the complex interplay of genes, proteins, cellular processes, and environmental factors from raw omics data. To facilitate the reproducibility of omics studies and the effective secondary analysis of omics data, metadata completeness and accuracy are crucial^4,5^. Metadata enriches raw data by providing essential details about its fundamental attributes, including phenotype, age, sex, and disease condition, as well as comprehensive experimental and environmental information, such as the data generator, creation date, data format, and sequencing protocols^2^. As a result, accurate and comprehensive metadata are critical for the efficient utilization, sharing, and subsequent re-analysis of omics data^1,6–10^. Metadata also enable data-driven decision-making and policy development in fields such as biomedical sciences, clinical research, environmental sciences, and social sciences^1^.

Although the biomedical community has made tremendous efforts in sharing omics data, little attention is allocated to ensure the completeness of metadata accompanying raw omics data^1,11^. We previously reported the limited availability of metadata accompanying sepsis-based transcriptomics studies^1^ but the overall patterns of metadata sharing across other diseases and organisms remains unknown^1,12,13^. Incomplete metadata hinders researchers’ ability to utilize metadata information for subsequent downstream analysis^8,14,15^. For instance, studies have shown that metadata elements such as sample source, sex, or experimental condition are often missing or inconsistently reported in public repositories like GEO and SRA, limiting reusability^1,16^. Even when raw data are deposited, the lack of standardized descriptors and structured fields impedes automated retrieval and integration^17^. In addition to hindering reproducibility, incomplete metadata can lead to underutilization of publicly funded datasets. A study analyzing metadata quality in gene expression repositories found that many samples lacked even basic information such as organism or tissue type, making them effectively unusable for secondary analyses or meta-studies^18^. This not only limits the scientific return on investment but also impedes large-scale integrative research that relies on well-annotated datasets. Typically, metadata accompanying omics studies has been shared in two ways, in public repositories and in the text of publications. Metadata shared solely in the textual content of publications has many limitations and is insufficient to ensure that it is complete, accurate, accessible, and machine-actionable^19^. This is because metadata not in a standardized format in publications can be scattered across multiple locations, making it challenging and time consuming for researchers to locate and integrate metadata from multiple sources for further downstream analysis. Additionally, metadata shared in publications is often provided at the study level lacking per sample or individual information. The lack of sample-level metadata limits the reproducibility of conducted research, restricting the credibility and ultimately making secondary analysis impossible or extremely difficult to conduct. If the metadata is shared only in the publications, the researchers will require a time-consuming and laborious manual data-mining process to extract such metadata information, especially when looking at large scale studies^20^. Mining and extracting metadata from the publications can be challenging due to the presence of misannotated, unstructured and absent metadata information. In contrast to sharing metadata within the text of a publication, a highly effective method for disseminating metadata involves sharing it through publicly accessible repositories, making researchers easy to access this information and facilitate downstream secondary analyses^1,21^. As a result, public repositories play a crucial role in advancing biomedical research by facilitating the efficient and effective sharing and utilization of data accompanying metadata^1^.

In this study, we analyzed a total of 253 randomly selected studies over 164 thousands samples across various disease conditions, phenotypes and organisms. We investigated the prevailing practice of metadata sharing in the textual content of publications and the corresponding public repositories. We observed that studies often omit over a quarter of crucial phenotypes, with an average of only 74.8% shared. Only 11.5% of studies completely shared all phenotypes, while 37.9% shared less than 40% of the phenotypes. Studies involving non-human samples were more likely to have complete metadata than studies involving human samples. Additionally, public repositories contained more complete metadata compared to metadata shared in the textual content of publications. To generalize our results, we further examined over 61 thousand studies over 2.1 million samples from Gene Expression Omnibus^22^ (GEO) repository. The overall metadata availability of 2.1 million GEO samples was 63.2%. Similar to the metadata availability of surveyed 253 studies, non-human studies have 16.1% more pheenotypes shared compared to human studies. Notably, the availability of metadata in published studies has increased substantially over time. In studies published before 2011, metadata availability was limited, with less than 1% of studies having metadata being shared. In contrast, in studies released after 2021, there has been a remarkable enhancement in metadata availability, with as many as 50% of studies now incorporating this valuable data.

Rich, machine-readable phenotypic metadata are now recognised as a prerequisite for transparency, reproducibility, and equitable reuse of biomedical omics data. International frameworks—most notably the FAIR Guiding Principles and the 2023 NIH Data Management and Sharing (DMS) policy—explicitly mandate that publicly shared datasets be accompanied by sufficiently detailed descriptors to enable findability and rigorous downstream analysis^23^. Yet community audits continue to show that key variables are routinely omitted, constraining reliable population stratification, sample matching, and mechanistic inference. To interrogate this gap we operationally define metadata completeness as the presence, at minimum, of six phenotypic attributes that recur across legacy and emerging minimum-information standards (MIAME^24^, MINSEQE^25^, MIBBI^26^, GA4GH Phenopackets^27^) and that exert direct analytical influence: race/ethnicity/ancestry (REA), age, sex, tissue type (or specific cell type), organism, and experimental strain information. These fields are mandated or strongly recommended in multiple checklists precisely because they capture the principal axes of biological heterogeneity that confound multi-study integration and drive disease risk. Attributes such as marital status or geographic latitude, while relevant in certain contexts, lack comparable consensus and are inconsistently reported; they were therefore excluded from our baseline definition.

Using this schema we examined 61,312 genomics and transcriptomics records deposited between 2008 and 2024. Automated XML parsing was complemented by a manual audit of 253 randomly selected studies, confirming parser accuracy and revealing silent discrepancies between manuscript narratives and repository entries. Fewer than one-third of studies reported all six attributes in either venue, and 42 % of the attributes that were disclosed in one source were absent from its counterpart. Although reporting rates have improved modestly since 2018—coincident with the roll-out of FAIR and NIH DMS guidance—completeness remains highly variable across sub-disciplines. Collectively, these findings substantiate our underlying assumption that the six selected attributes constitute a practical yet stringent benchmark for evaluating metadata sufficiency, and they underscore the urgent need for harmonised publisher–repository checklists, enforceable submission validators, and community incentives that treat metadata as a first-class research output. Addressing these structural barriers will accelerate hypothesis generation, cross-study integration, and ultimately clinical translation across the biomedical research spectrum. By promoting the availability and quality of metadata accompanying raw omics data, we can enable more accurate and efficient secondary analyses, which in turn may advance our understanding of complex diseases and their underlying mechanisms, and ultimately discover novel biomedical insights to improve human health^28^.

## Results

### Assessing the completeness of public metadata accompanying omics studies

We have performed a comprehensive analysis of the availability of the metadata reported in the textual content of the original scientific publications and the corresponding public repositories. We meticulously analyzed a total of 253 randomly selected scientific publications encompassing over 164 thousand samples across various disease conditions, phenotypes and organisms (D1 dataset) (Supplementary table 1). The human studies encompassed various disease conditions, namely Alzheimer’s disease (AD), acute myeloid leukemia (AML), cardiovascular disease (CVD), inflammatory bowel disease (IBD), multiple sclerosis (MS), sepsis, and tuberculosis (TB) (Table S1) (Figure S1). Among the 253 studies, 153 pertained to human studies, while 100 focused on non-human studies. To assess metadata availability, we manually reviewed the phenotypic information within the textual content of publications, including both the main text and supplementary materials. To retrieve metadata from the public repositories, we developed the custom Python scripts (Methods Section). To increase the generalizability of our analysis, we randomly selected over 61 thousands studies from the Gene Expression Omnibus^29^ (GEO) repository, encompassing a total of over 2.1 million samples (D2 dataset) (Supplementary table 2). For both D1 and D2 datasets, we examined the four common phenotypes shared by both human and non-human studies, which included organism, sex, age, and tissue types. In addition to the common phenotypes, we investigated the human-specific phenotype of race/ethnicity/ancestry and the non-human-specific phenotype of strain information.

### Limited availability for essential phenotypes accompanying 253 studies

Our analysis has unveiled that more than a quarter of phenotypes are not shared in either the textual context of the publication or public repositories (Figure S2). While all studies shared at least one phenotype, on average, only 11.5% of studies managed to encompass the entire set of phenotypes. In contrast, over a quarter of studies shared a significant 80% of the phenotypes, and more than a third of studies shared 40% of the phenotypes. Additionally, 4.7% of the studies shared less than 20% of the phenotypes (Figure 1a). Among the six phenotypes, organism information was the most commonly shared phenotype with all the studies sharing such information (Figure 1b). Meanwhile, up to 80% of the studies reported tissue information, and similarly, 80% of the non-human samples reported strain information (Figure 1b). About 60% of the studies have available sex and age information (Figure 1b), with, on average, half of the samples within the studies reporting this information in both sources. Importantly, the race/ethnicity/ancestry information was the least frequently reported phenotype with only up to 20% of the human studies reported this essential phenotype information (Figure 1b), with only about 10% of the samples, on average, sharing such information within the studies in both sources. Our analysis revealed a substantial improvement in metadata reporting practices, particularly in studies released after 2018 (Figure 1c). Most of the phenotypes experienced a drastic increase in availability after 2018, including age, organism, sex, tissue, and strain information. In contrast, studies published before 2010 show a significant limitation in the availability of the six phenotypes (Figure S3).

**Figure 1.**
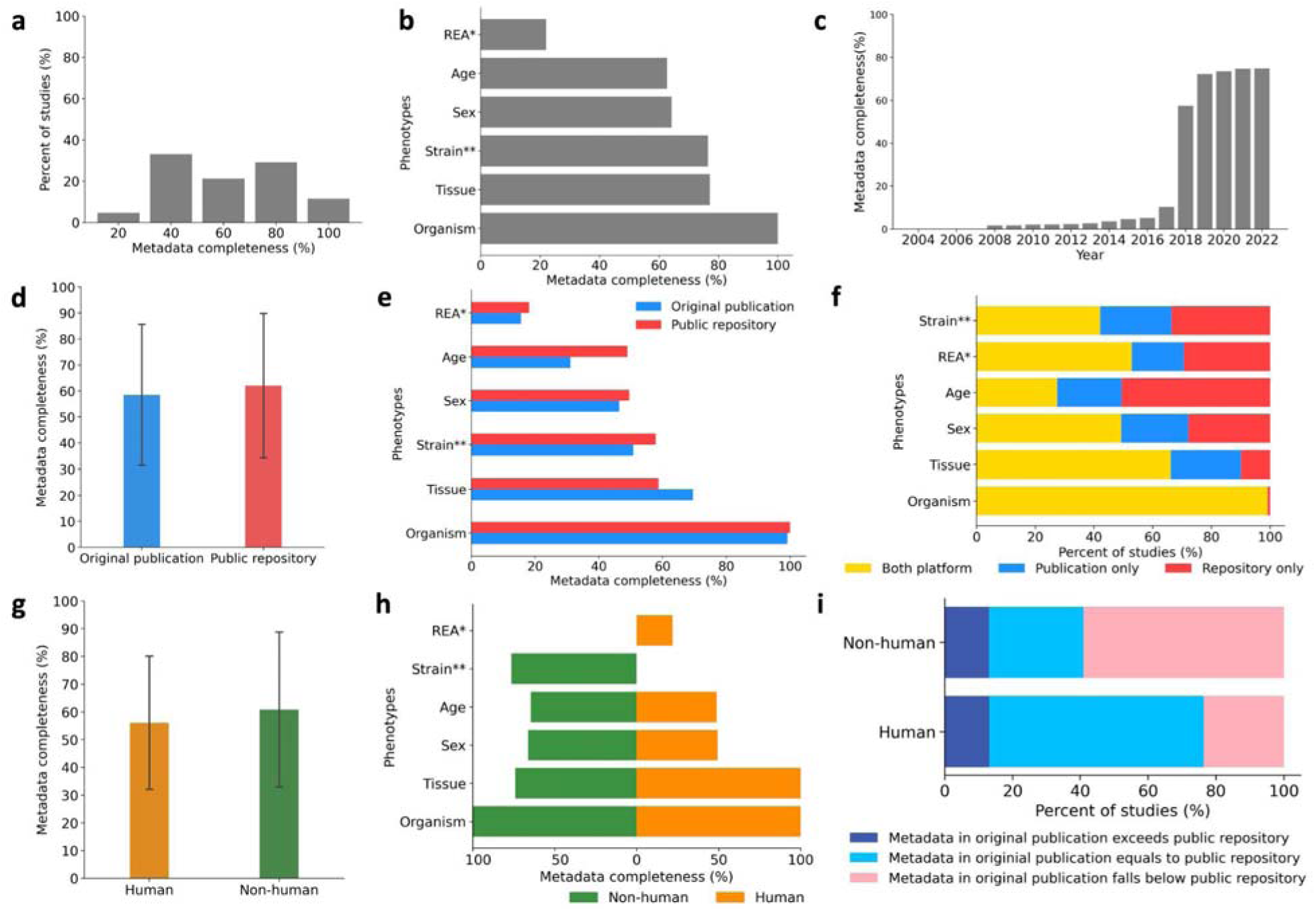
Completeness of public metadata accompanying omics studies (a) The distribution of studies that share 100% of metadata (all phenotypes), 80% of metadata (four phenotypes), 60% of metadata (three phenotypes), 40% of metadata (two phenotypes), and 20% of metadata (one phenotype) in the D1 dataset. (b) The availability of the six phenotypes information in the textual content of publications and/or public repositories in the D1-dataset. (c) The cumulative metadata availability of the D1-dataset over the years. (d) The overall metadata availability in the textual content of publications and public repositories in the D1-dataset. (e) The sample-level metadata across seven phenotypes among all the samples in the D1-dataset reported in the textual content of publications and public repositories. (f) The proportion between the studies shared the six phenotypes in both sources, including textual content of the publication and public repositories, the studies shared the six phenotypes only in the textual content of the publication, and the studies shared the six phenotypes only in the public repositories. (g) The overall metadata availability for human and non-human samples in the D1-dataset. (h) Right: The overall composition of sample-level metadata availability across six phenotypes in the textual content of publications and public repositories across the human samples in the D1- dataset; Left: The overall composition of sample-level metadata availability across five phenotypes in the textual content of publications and public repositories across the non-human samples in the D1-dataset. (i) The percentage of non-human and human studies in the D1-dataset that reported the sample-level metadata in the textual content of publications the same as, exceeds, or falls below the sample-level metadata reported in the public repository. (The REA* (race/ethnicity/ancestry) information is analyzed over the human samples only; The strain** information is analyzed over the non-human samples only.)

### Inconsistent practices of metadata sharing across the textual content of publications and corresponding public omics repositories

We compared metadata availability across the 253 studies reported in the textual content of publications versus in the corresponding public repositories. The metadata availability in original publications was more than 3% lower than the metadata availability in public repositories, which was 62.0% (Figure 1d). In comparison to the textual context of the publications, public repositories share more complete phenotypes for organism, sex, age, strain, and race/ethnicity/ancestry, while the textual content of publications share more complete tissue information (Figure 1e). Organism information was the most completely and consistently shared phenotype in both public repositories and the textual content of the publications with all such information available across all the 164,909 samples (Figure 1e). Notably, organism information was fully available in the public repositories in contrast to 99.2% availability in the textual content of the publication (Figure 1e). In contrast, age was the least consistently shared phenotype between the textual content of publications and public repositories, with 17.9% more samples having age information in public repositories (Figure 1e). Tissue information was reported in both platforms for over 60% of the samples, 20% of samples only report such information in the textual content of the publications, while only about 10% shared it only in public repositories. About 50% of the samples shared sex and race/ethnicity/ancestry information between both sources, and 30% of the samples only shared such information in public repositories (Figure 1f).

### Non-human studies demonstrated an enhanced commitment to sharing metadata

We compared the metadata reporting practices for the common phenotypes across 153 non-human and 100 human studies. The metadata availability for human samples was 56.1% and for non-human samples was 60.86% (Figure 1g). Human and non-human samples had more complete metadata reported in public repositories (66.6% and 62.0%, respectively) than in the textual content of publications (54.6% and 59.7%, respectively) (Figure S4). Non-human studies reported more complete metadata for age and sex, while human studies reported more complete metadata for tissue information (Figure 1h). In the human and non-human studies, organism information is widely available (Figure 1h). Tissue information is also widely available in human studies; conversely, only up to 60% of the non-human samples had available tissue information. While organism and tissue information was commonly reported in human studies, sex data was provided by only 49.4% of human studies, age data by 48.9%, and race/ethnicity/ancestry details by a mere 22% of human studies (Figure 1h). For non-human studies, other than the organism data, strain information was the most reported, available for 76.5% of samples. Age, and sex information has over 60% availability in non-human studies (Figure 1h). We observed that both human and non-human studies shared more complete metadata in public repositories than in the textual content of publications. Of the 153 human studies, 97 had consistent metadata sharing practices between original publications and public repositories, 36 had lower metadata availability in the textual content of publications, and 20 had higher metadata availability in the textual content of publications (Figure 1i). Of the 100 non-human studies, 59 had more complete metadata in public repositories than in the textual content of publications, 28 had the same extent of metadata availability in both sources, and 13 had more complete metadata in the textual content of publications than in public repositories (Figure 1i).

We observed significant discrepancies in metadata sharing among non-human species and aim to assess the overall completeness of metadata across these species. We analyzed metadata availability in the textual content of publications and public repositories for the top five non-human organisms in the 153 non-human studies, including Mus musculus, Parus major, Glycine max, Danio rerio, and Gallus gallus. Mus musculus and Parus major had similar metadata availability levels (up to 60%) in both sources. In contrast, Glycine max, Danio rerio, and Gallus gallus demonstrate a notable difference in metadata availability between the textual content of publications and public repositories. Specifically, they reported more comprehensive metadata in the textual content of publications than in public repositories (Figure S5).

### Sharing experiment-level metadata remains a common practice

The concept of experiment-level metadata availability was characterized by the sharing of only summarized information regarding the phenotypes of the samples, omitting detailed per-sample phenotypic information. It remains a common practice to offer an overall description of the study or experiment’s participants, while refraining from providing detailed descriptions of each individual’s phenotypes. Among human studies, 14.8% of metadata was exclusively shared at the experiment-level, while non-human studies exhibited a comparable pattern, with 6.8% of metadata exclusively shared at the experiment-level. In human studies, approximately 60% of the samples reported available age information only at the experiment-level (Figure S6). Up to 20% of the human samples only shared experiment-level metadata for disease, sex, and race/ethnicity/ancestry information (Figure S6 and Figure S7). In non-human studies, 20% of the non-human samples only reported tissue information at the experiment-level. Sex information was often presented in a summarized format, with approximately 40% of this information available at the experiment level. In contrast, less than 10% of the sex information was available at the sample level (Figure S6 and Figure S7). Notably, the experiment-level metadata for sex, age, and race/ethnicity/ancestry information accounts for approximately half of the human samples’ phenotypic information availability (Figure S7).

### Discrepancies in metadata sharing practices were observed across seven disease conditions Next, we examined metadata sharing practices for seven disease conditions, including

Alzheimer’s disease (AD), acute myeloid leukemia (AML), cardiovascular disease (CVD), inflammatory bowel disease (IBD), multiple sclerosis (MS), sepsis, and tuberculosis (TB), in human studies. The metadata completeness for the six phenotypes varies across the seven disease condition studies (Figure 2). All seven diseases have fully reported metadata for organism and tissue types (Figure 2). The reporting practices of three phenotypes (sex, age, and race/ethnicity/ancestry information) exhibited variability across the seven diseases. For the sex information, CVD studies had reported the highest availability for such information (75.9%), followed by AD studies, sepsis studies, AML studies, IBD studies, and TB studies, while MS studies had the lowest (10.8%). For the age information, sepsis studies had reported the highest availability of such information (67.9%), followed by AD studies, AML studies, IBD studies, CVD studies, and TB studies (32.4%), while MS studies had the lowest (9.7%). For the race/ethnicity/ancestry information, most studies rarely reported such information, and notably, none of the samples in the IBD and MS studies reported race/ethnicity/ancestry information (Figure 2).

**Figure 2.**
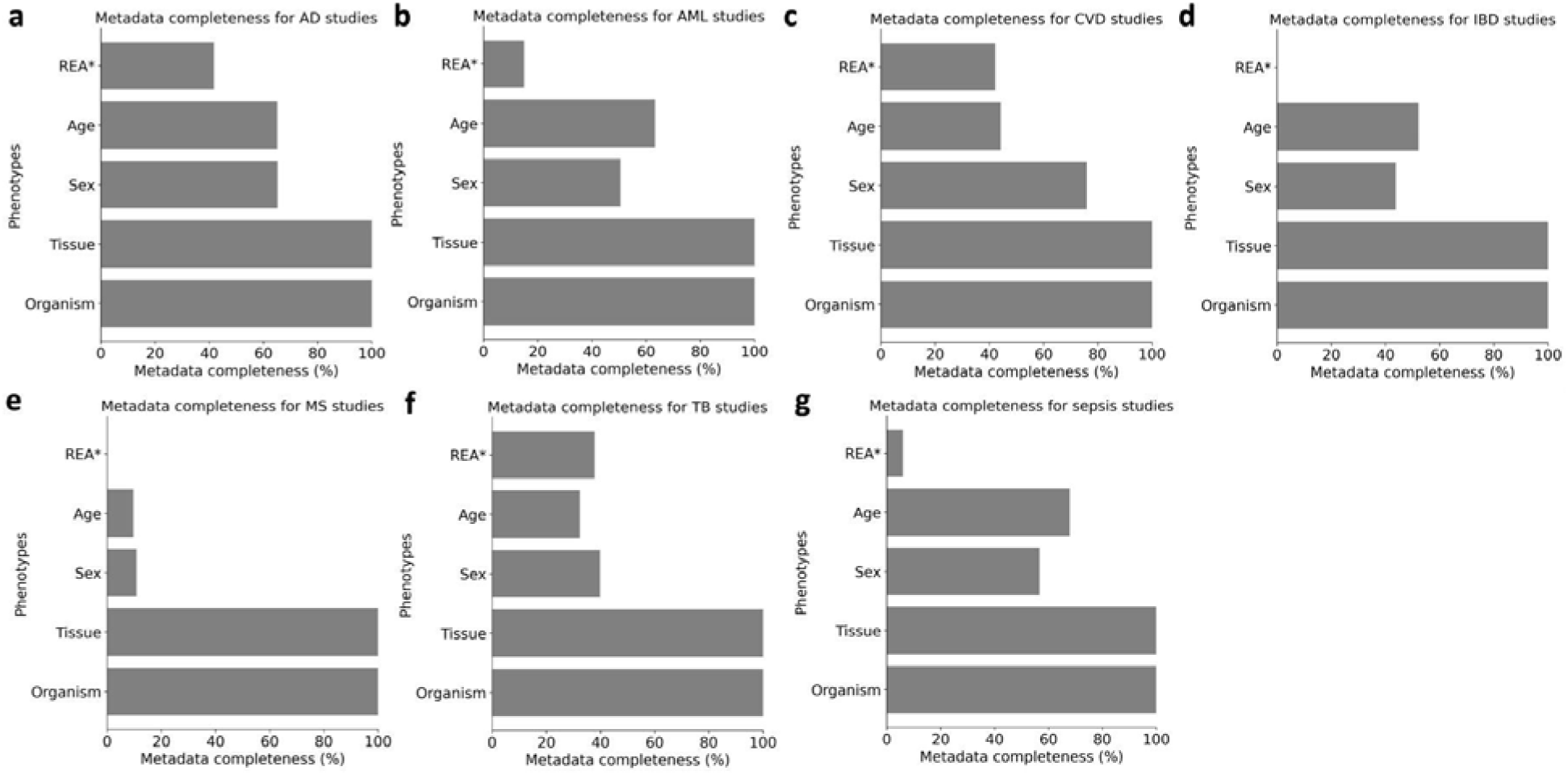
Metadata completeness across human studies. (a). Metadata completeness for Alzheimer’s disease (AD) studies. (b) Metadata completeness for acute myeloid leukemia (AML) studies. (c) Metadata completeness for cardiovascular disease (CVD) studies. (d) Metadata completeness for inflammatory bowel disease (IBD) studies. (e) Metadata completeness for multiple sclerosis (MS) studies. (f) Metadata completeness for tuberculosis (TB) studies. (g) Metadata completeness for sepsis studies. (The REA* (race/ethnicity/ancestry) information is analyzed over the human samples only; The strain** information is analyzed over the non-human samples only.)

Analysis of 61,000 studies encompassing 2.1 million samples across public repositories confirms limited metadata availability

We investigated the availability of the six phenotypes reported for 2,168,620 samples from 61,950 studies available at the Gene Expression Omnibus (GEO) (D2 dataset). The overall availability of metadata among the 61,950 studies was 63.2% (Figure S8). Notably, we discovered that studies preceding 2010 exhibited a lack of shared metadata (Figure S9). We discovered that human studies exhibited a lower average metadata completeness (47.4%) compared to non-human studies (63.5%) (Figure S8). As a result, the availability of metadata in non-human samples surpasses that in human samples (Figure S8 and Figure S9). Human samples invariably share organism and tissue information (100% and 99.9%, respectively), but less frequently report age (19%) and sex (13.8%), with race being the least disclosed at 4.2% (Figure S10); Conversely, all non-human samples report organism and tissue data, with strain and age information present in 56.7% and 37.4% of samples, respectively, while only 24.15% include sex information (Figure S10). Tissue, sex, and age metadata are more comprehensively reported in non-human samples compared to their human counterparts (Figure S10).

## Discussion

In our study, we focused on six phenotypic attributes—organism, tissue type, strain, sex, age, and race/ethnicity/ancestry (REA)—as representative indicators of metadata completeness. This selection was guided by a combination of field-wide reporting standards and their relevance to downstream analyses. We used standardized descriptors—such as organism, tissue or cell type, and experimental strain—guided by international minimum-information checklists (e.g., MIAME^24^, MINSEQE^25^, MIBBI^26^, GA4GH Phenopackets^27^), as these phenotypes are consistently reported across studies and essential for selecting appropriate reference genomes, expression atlases, and functional annotations. Their omission has been shown to inflate false-positive rates in differential-expression pipelines when samples from distinct tissues or mouse substrains are inadvertently pooled. In particular, strain information in non-human model organisms (e.g., mice, rats, zebrafish) is essential for reproducibility in molecular studies, especially in areas such as epigenomics, where strain-specific genetic backgrounds can significantly influence DNA methylation patterns, histone modifications, and chromatin accessibility. For example, studies have shown that genetically similar mouse strains exhibit distinct epigenetic profiles that affect disease susceptibility, gene regulation, and developmental phenotypes. The absence of such metadata can lead to misinterpretation of epigenetic variation as disease-related rather than background-driven^30,31^.

Certain phenotypic variables—such as sex, age, and ancestry—carry substantial biological and analytical value, and their inclusion is essential for accurate, reproducible, and equitable interpretation of genomic data. The NIH “Sex as a Biological Variable” (SABV) policy (NOT-OD-15-102) codifies sex as essential for reproducibility; meta-analyses of pre- and post-policy publications show that failure to stratify by sex conceals up to 22 % of significant expression differences^32–34^. As a result, sex is widely recognized as a critical biological variable; it has been strongly recommended by the NIH and others as essential for identifying sex-specific disease mechanisms, treatment responses, and biological variability^35^. Age profoundly modulates transcript abundance (e.g., >40 % of protein-coding genes in GTEx exhibit age-dependent expression), alters epigenetic drift, and drives batch effects in single-cell atlases; downstream analyses that ignore age confounders mis-estimate effect sizes and disease signatures^36–38^. Race/Ethnicity/Ancestry (REA) captures population-specific allele frequencies and gene-environment interactions. Large-scale surveys consistently show that >80 % of GWAS participants are of European ancestry; re-analyses demonstrate that polygenic risk scores trained on such data lose predictive power in under-represented groups, underscoring the necessity of REA metadata for equitable translational research^39^. Recent studies have also emphasized that the lack of ancestral diversity in genomics undermines generalizability and equity in data-driven discoveries^40,41^.

Collectively, these six attributes represent the intersection of (i) widespread standardisation, (ii) availability across public repositories, and (iii) demonstrable influence on downstream bioinformatic pipelines. Their absence hampers sample matching, inflates hidden confounders, and ultimately erodes reproducibility—as evidenced by the replication failures cited above. While additional clinical variables (e.g., disease stage) are invaluable in specific contexts, they are neither universally reported nor mandated by current standards, precluding systematic cross-study assessment at present. Our definition of metadata completeness therefore reflects a pragmatic yet stringent baseline that is both measurable and maximally informative across heterogeneous study designs.

Although additional metadata elements—such as disease severity, batch, or sampling location—can be valuable in specific study contexts, the six we selected are expected to be routinely collected and are broadly informative across study designs, even if not always publicly available. The absence of these variables can limit sample stratification, introduce confounding, and reduce the ability to replicate study findings—thereby undermining reproducibility and interpretability in secondary analyses. Our analysis emphasized phenotypes that are broadly available and comparable across a wide range of study types. While clinical metadata—such as treatment history, disease severity, or comorbidities—are essential in clinically focused research, such information is often absent in basic or translational studies that fall under disease categories but focus primarily on molecular or mechanistic biology. In these contexts, clinical metadata may not be reported or applicable, making systematic extraction and cross-study comparison challenging. We acknowledge the importance of assessing clinical phenotype availability and view this as a valuable direction for future research, particularly in studies that target clinical datasets.

Our study is the first to systematically analyze metadata completeness across multiple organisms and phenotypes, revealing limited availability of metadata both in textual content of publications and public repositories. While metadata completeness varies by phenotype, organism and tissue types are consistently reported, whereas age, sex, strain, and race/ethnicity/ancestry information are often missing (Figure S2). Notably, non-human studies demonstrated an enhanced commitment to sharing metadata than human studies. Several factors may contribute to the observed differences in metadata availability between human and non-human studies. Firstly, the nature of the research focus plays a crucial role. Human studies are typically subject to stricter ethical guidelines and consent requirements, and data sharing regulations^42–44^. These regulations may limit the extent of metadata that can be collected, whereas non-human studies may not encounter the same constraints^42,44–46^. As a result, non-human studies may enjoy easier access to metadata due to less stringent privacy and consent regulations, facilitating more extensive data collection efforts^47–49^. Data sharing norms may further influence metadata availability. Non-human studies often have a stronger tradition of data sharing, motivating researchers to provide more complete metadata to promote transparency and collaboration^50^.

Different research fields have specific requirements and conventions for metadata, which can explain the varying availability between human and non-human species. The nature and objectives of a study often dictate which metadata elements are deemed essential. For example, in disease model research—particularly cancer studies using mouse models^51^— detailed metadata such as sex, age, genetic background, and strain are often critical, as these variables can substantially influence phenotypic outcomes, tumor progression, and treatment response. In some studies, sex is not only a biological descriptor but also a key experimental variable, prompting researchers to prioritize and report such information comprehensively to ensure scientific rigor, reproducibility, and translational relevance^52^. Conversely, studies situated within evolutionary biology or population genetics tend to emphasize broader-scale attributes, such as species classification, sampling location, or population structure, while individual-specific metadata (e.g., age, sex, strain) may be of secondary importance or entirely absent. This difference in research focus leads to variable metadata completeness across species and study types. Additionally, standardization and reporting practices differ between research communities: fields with mature metadata standards and well-established reporting frameworks (e.g., biomedical mouse model research) typically exhibit greater metadata completeness, whereas emerging or interdisciplinary areas may lack uniform metadata protocols, resulting in inconsistencies^53^. Repository policies also play a role. Some genomic repositories enforce strict metadata submission guidelines for certain study types or species, while others offer more flexibility, contributing to the heterogeneity in metadata quality. Taken together, these contextual and infrastructural factors help explain why metadata reporting practices vary substantially across non-human studies and reinforce the importance of tailoring metadata standards to the needs of specific research communities.

Our study is also the first to investigate metadata availability at both the experiment and sample levels. Our previous analysis did not distinguish among the various methods of metadata sharing, whether it involved sharing metadata information for each sample individually or just providing summary statistics for the entire study^1^. Typically, sample-level metadata is more valuable for ensuring the transparency and reproducibility of reported results and for secondary analysis than experiment-level metadata, because it provides detailed information about each sample rather than a summary of the entire study^54^. Researchers who want to replicate or expand on an experiment or perform secondary analyses on the raw data need sample-level metadata^55^. In contrast, experiment-level metadata provides a comprehensive description of the study, but it may not be enough for many types of analyses that require a granular level of detail. When conducting analyses for new research questions, the specificity of sample-level metadata enables researchers to reuse the existing dataset to answer novel research questions and draw meaningful conclusions. We observed that metadata is often reported at the experiment-level, not the sample-level. In most textual content of publications, metadata is only available at the experiment level. We found that half of the human samples with available metadata for sex, age, and race/ethnicity/ancestry have only experiment-level metadata. Note that all studies have shared a portion of information pertaining to the metadata of the samples, implying a conscientious effort by the authors to provide relevant details about the samples’ metadata; however, such effort remains incomplete. However, there are several limitations in our study. First, we were able to only extract commonly reported phenotypes and were unable to assess the availability of study-specific phenotypes. Additionally, our analysis focused solely on the availability of metadata reported and did not address the quality or accuracy of metadata reported.

Making metadata widely available in public repositories such as GEO can enhance the transparency and reproducibility of research, facilitate cross-study comparisons, and contribute to the advancement of scientific knowledge by providing comprehensive contextual information about datasets. To ensure the quality and usefulness of clinical data, metadata reporting in all categories must be improved and shared across domains on public repositories, in addition to the text of publications^1^. Additionally, sharing metadata solely within the text of a publication is not an optimal approach for metadata sharing for several reasons. Firstly, when metadata is embedded within textual content, it becomes a challenge to locate and extract, especially for those researchers who may be sourcing multiple papers for comprehensive data. Additionally, the metadata presented within a publication might not always be complete; key details might be overlooked or omitted due to space constraints or editorial decisions. This potential for incompleteness can lead to inaccuracies when researchers rely on this data for their work. Moreover, when metadata is constrained within a publication, it’s not easily shareable. Digital repositories or databases, on the other hand, allow for streamlined sharing and collaboration. Thus, relegating metadata solely to the text of a publication creates barriers to efficient research and collaboration. Sharing metadata on public repositories can reduce the efforts in metadata mining directly from the textual content of publications or from the request to the authors since the both processes can be time-consuming and error-prone^11,56–58^. Leveraging previously published data for novel biological discoveries could be facilitated when the metadata accompanying its raw omics data is reported, present in a standardized format, and made available in online repositories^5,59,60^. Improving the availability of metadata in public repositories could provide valuable and accurate information for downstream analyses, and further enhance the usefulness of the public repositories.

To address the observed gaps in metadata availability, we outline several actionable strategies for improving reporting practices across the community. To improve metadata reporting in omics research, we propose four key strategies centered on standardization, automation, enforcement, and community coordination. First, adopting standardized metadata schemas based on the FAIR (Findable, Accessible, Interoperable, Reusable)^23,53,61^ principles is essential for ensuring data reusability and interoperability. Frameworks^62^ such as MIxS from the Genomic Standards Consortium^63^, GA4GH models^64^, and Clinical Data Interchange Standards Consortium^65^ (CDISC) templates provide structured guidelines for genomic and clinical metadata reporting. By adopting these frameworks, researchers can ensure that their metadata is comprehensive and consistent, thereby enhancing data integration and interoperability. Second, integrating automated tools can enhance metadata accuracy and reduce manual effort. Tools like Omics Metadata Management Software^66^ (OMMS) and MARMoSET^67^ enable standardized and automated metadata curation for various omics datasets, streamlining data preparation for analysis and sharing. Third, journals and funding agencies should mandate the inclusion of comprehensive metadata using standardized checklists aligned with FAIR principles. This requirement would promote transparency and reproducibility across research outputs. For instance, the Digital Repository of Ireland^68^ (DRI) emphasizes the importance of adhering to the FAIR principles to enhance data sharing and reuse. By enforcing such standards, journals and funding bodies can drive cultural change towards more transparent and reproducible research practices. Finally, establishing community-wide minimal metadata reporting guidelines tailored to specific omics data types can reduce ambiguity and improve data integration. Efforts led by groups like the Genomic Standards Consortium (GSC) have demonstrated the value of domain-specific consensus standards^23,62,69^. By implementing these strategies, the scientific community can address current challenges in metadata reporting, thereby enhancing the reproducibility, reliability, and collaborative potential of omics research^61^.

Our overall findings highlight the need for improved metadata reporting and sharing practices in biomedical research. The limited availability of metadata in both textual content of publications and public repositories impedes data reuse and reproducibility, and the disparities in completeness across different phenotypes make it difficult to conduct secondary analyses. We encourage authors to routinely report all relevant metadata, including organism, tissue type, age, sex, strain, and race/ethnicity/ancestry information, in both their publications and public repositories. Improving the accessibility and reliability of metadata would significantly benefit the broader scientific community by facilitating data-driven research and fostering secondary analysis within the biomedical research field.

Our metadata extraction strategy focused on structured MINiML XML records available from the Gene Expression Omnibus (GEO) via the NCBI FTP server. These XML files were selected because they represent the only uniformly formatted, machine-readable metadata source consistently provided for every GEO study. Their standardized schema and widespread accessibility make them a pragmatic choice for cross-study and cross-domain assessments of metadata completeness. Prior studies have shown that structured formats like XML enhance transparency and reusability, especially when accompanied by schema definitions and parsing tools^70^. To evaluate the accuracy of our XML-based extraction pipeline, we manually reviewed all 253 studies in our curated D1 subset. This included studies labeled by the pipeline as missing one or more of the six core attributes. In each case, we confirmed that the XML output accurately reflected the metadata reported in the GEO record and associated publication materials. This validation provides strong support for the fidelity of our automated approach, particularly in large-scale applications where manual inspection is infeasible. Nonetheless, we acknowledge that metadata stored exclusively in alternative formats—such as HDF5 (.h5ad), R data files (.rds), spreadsheets, or supplemental PDFs—was not captured by our pipeline. These formats are often fragmented, lack standardization, or are embedded within proprietary analysis environments, limiting their accessibility to non-specialist users. While our findings reflect the most structured and broadly accessible metadata available in GEO, this design choice may lead to a modest underestimation of overall metadata availability—especially in rapidly evolving areas such as single-cell omics. Future efforts may benefit from extending metadata extraction tools to accommodate these emerging formats.

Moreover, metadata reporting across studies is often inconsistent, varying in terminology, structure, and completeness. These inconsistencies introduce challenges in automated parsing and can impact the comparability of results across datasets. For example, key attributes such as race/ethnicity/ancestry (REA) and tissue type are frequently labeled in non-standardized ways, making them difficult to detect reliably. While our study focused on assessing the prevalence of reported metadata, a comprehensive evaluation of the quality, accuracy, and downstream scientific impact of missing metadata is outside the current scope.

To partially mitigate these limitations, we conducted a detailed manual review of 253 studies to evaluate metadata availability beyond what could be captured automatically. However, we acknowledge that even this curated subset may not fully capture metadata provided exclusively in complex or proprietary formats. Moving forward, the development of more adaptable, format-aware extraction tools and benchmarking against manually annotated corpora will be essential for improving both the accuracy and comprehensiveness of metadata completeness assessments in large-scale omics repositories.

Several developments around 2018 may have contributed to improved metadata reporting practices observed in public repositories. For instance, many journals began adopting standardized data policies aimed at enhancing transparency and reproducibility. Springer Nature, for example, implemented standard data policies across over 1,500 journals, encouraging researchers to share data and associated metadata openly^71^. Concurrently, community-driven efforts such as Crossref’s reintroduction of Participation Reports helped promote open metadata standards by providing feedback on the completeness of metadata records. Moreover, the broader open science movement gained momentum during this period, with initiatives such as Plan S^72^ advocating for immediate open access to research outputs and reinforcing the importance of metadata for enabling data reuse. While these initiatives likely influenced the trends we observed, we acknowledge that our current study does not establish a direct causal relationship between policy changes and metadata completeness. Further investigation would be required to systematically assess the impact of these external factors.

Lastly, while the GEO constitutes a highly comprehensive repository for omics data and associated metadata, it represents only a singular component within the broader ecosystem of publicly accessible genomic repositories. Our analysis focused specifically on the GEO and consequently excluded other significant repositories such as the Sequence Read Archive (SRA), European Nucleotide Archive (ENA), ArrayExpress, or domain-specific databases. Future research endeavors should aim to broaden this analysis to encompass a more comprehensive suite of these resources. Such an expanded investigation would yield a more robust and representative assessment of metadata reporting standards and practices across the wider genomics research landscape.

## Methods

### Datasets

We examine the per-sample metadata completeness within the D1 dataset, spanning 253 mammalian multi-omics studies retrieved from NCBI’s Gene Expression Omnibus (GEO) and comprising a total of 164,909 samples. This dataset included 20,047 human samples from 153 studies and 144,862 non-human samples from 100 studies. The human studies covered a variety of diseases, including Alzheimer’s disease, acute myeloid leukemia, cardiovascular disease, inflammatory bowel disease, multiple sclerosis, sepsis, and tuberculosis. The remaining 100 studies involved non-human samples, including Mus musculus, Parus major, and Glycine max (Figure S1). Additionally, for the D2 dataset, we randomly selected 61,950 studies from the GEO repository, encompassing a total of 2,168,620 samples. Among the studies in the D2-dataset, 2,119,749 samples were non-human samples, while 48,871 samples were human samples.

### Examined phenotypes

We examined the metadata for the six phenotypes among the human and non-human studies in the D1 and D2 dataset. For human samples, we examined the human-specific phenotype of race/ethnicity/ancestry information, along with four common phenotypes, including organism, age, sex, and tissue types. Conversely, for non-human samples, we investigated the additional non-human-specific phenotype of strain information, along with the four common phenotypes, including organism, age, sex, and tissue types. In the D2 dataset, we examined the metadata for the same six phenotypes as in the D1 dataset, including tissue types, organism, sex, age, strain information (non-human samples), and race/ethnicity/ancestry (human samples).

### Extraction of metadata from the textual content of publications

We manually extracted metadata from the main text and supplementary materials of each publication. Based on the information provided, we classified metadata availability into two categories: sample-level metadata available and experiment-level metadata available. Sample-level metadata was categorized as available only if the study explicitly shared metadata for individual samples. Additionally, if the metadata was not explicitly provided for each sample but could be inferred with reasonable certainty (e.g., study says that all samples are female), we also classified it as sample-level metadata. In contrast, studies that provided only summary metadata without individual sample-level details were classified as experiment-level metadata available and they don’t indicate that all samples carry the same phenotype. These studies usually included summary metadata information that could not be linked to specific samples’ metadata information.

### Extraction of metadata from the public genomic repositories

#### Accession list generation

On 12 April 2025 we retrieved the GEO “series family” index and compiled 61 312 unique Series (GSE) accessions covering studies released between 2008 and 2024. For each accession we recorded the location of its MINiML archive on the NCBI FTP server, housed in the tranche directory corresponding to its numeric range (e.g. the archive for GSE123456 resides in the directory GSE123nnn).

#### Automated download

A custom Python workflow established anonymous FTP sessions and systematically downloaded every MINiML-formatted XML archive. Compressed files were decompressed in situ, and automatic retry logic resolved transient network interruptions. The final retrieval success rate exceeded 99.9 %.

XML parsing and metadata normalisation Each MINiML file was parsed and converted into structured tabular form. Two tables were produced per study:

- a Series-level table recording accession, title, submission date, release date and overall experimental design;
- a Sample-level table capturing six core phenotypic attributes—organism, tissue/cell type, strain, sex, age and race/ethnicity/ancestry (REA)—together with additional descriptors such as library strategy and sample description.

Free-text entries were harmonised by applying rule-based string matching and ontology look-ups against widely used vocabularies (NCBI Taxonomy, Uberon, Cell Ontology and HANCESTRO). Manual validation

To evaluate the accuracy of our metadata extraction pipeline, we conducted a manual review of all 253 studies in the curated validation subset (D1), including those flagged by the automated workflow as missing one or more core attributes. For each study, we cross-validated the extracted XML metadata against information available elsewhere in the corresponding GEO entry or associated publication. This audit revealed complete concordance (100%) between the automated extraction and manual inspection for all six core attributes.

#### Scope and limitations

The pipeline operates exclusively on the MINiML XML channel—the standardized, machine-readable metadata format provided for all GEO submissions. Phenotypic information available solely through other formats (e.g., HDF5, R data files, spreadsheets, or PDF supplements) is not captured. Manual validation of 253 studies suggests that such instances are rare and do not impact the prevalence trends reported here. However, this design choice may modestly underestimate absolute metadata completeness, as discussed in the Discussion section.

#### Contributions

SM conceived of the presented idea and supervised the work. YH analyzed the collected data and drafted the manuscript. PJ validated the collected data from human studies. AR collected the data from sepsis studies. RG and EL collected the data from tuberculosis studies. IN collected the data from cardiovascular disease studies. RA and MYW collected data from acute myeloid leukemia. JH collected data from inflammatory bowel disease studies. AS and AN collected data from Alzheimer’s disease studies. SK collected data from non-human studies. YH, and GB developed customized Python scripts to extract metadata from GEO. All co-authors contributed to the revision of the manuscript.

## Code Availability

All code for the study can be found at https://github.com/Mangul-Lab-USC/metadata-completeness.

## Data Availability

All data for the study can be found at https://github.com/Mangul-Lab-USC/metadata-completeness and the supplementary materials.

## Supplementary Figures

**Figure S1.**
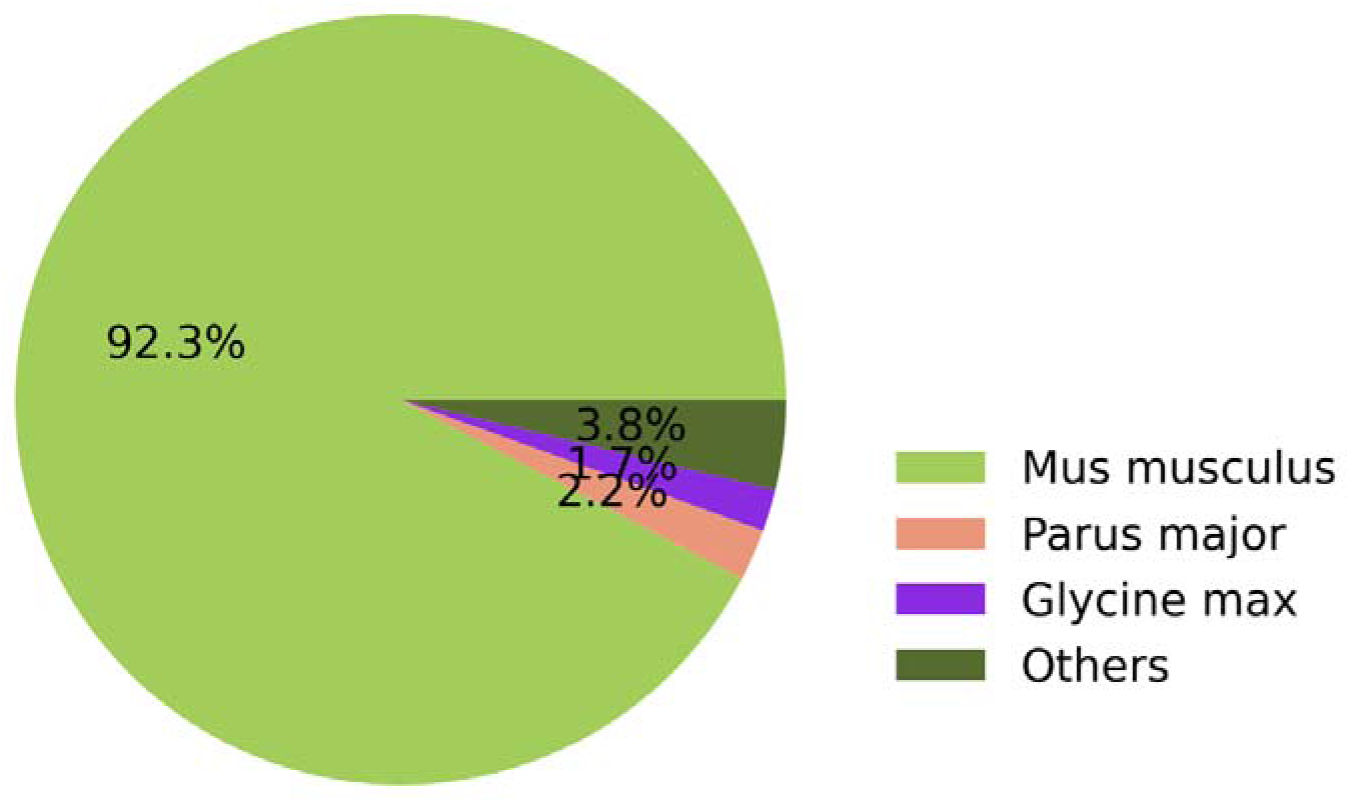
The proportion of non-human samples across 100 studies encompassing 144,862 samples in the D1 dataset.

**Figure S2.**
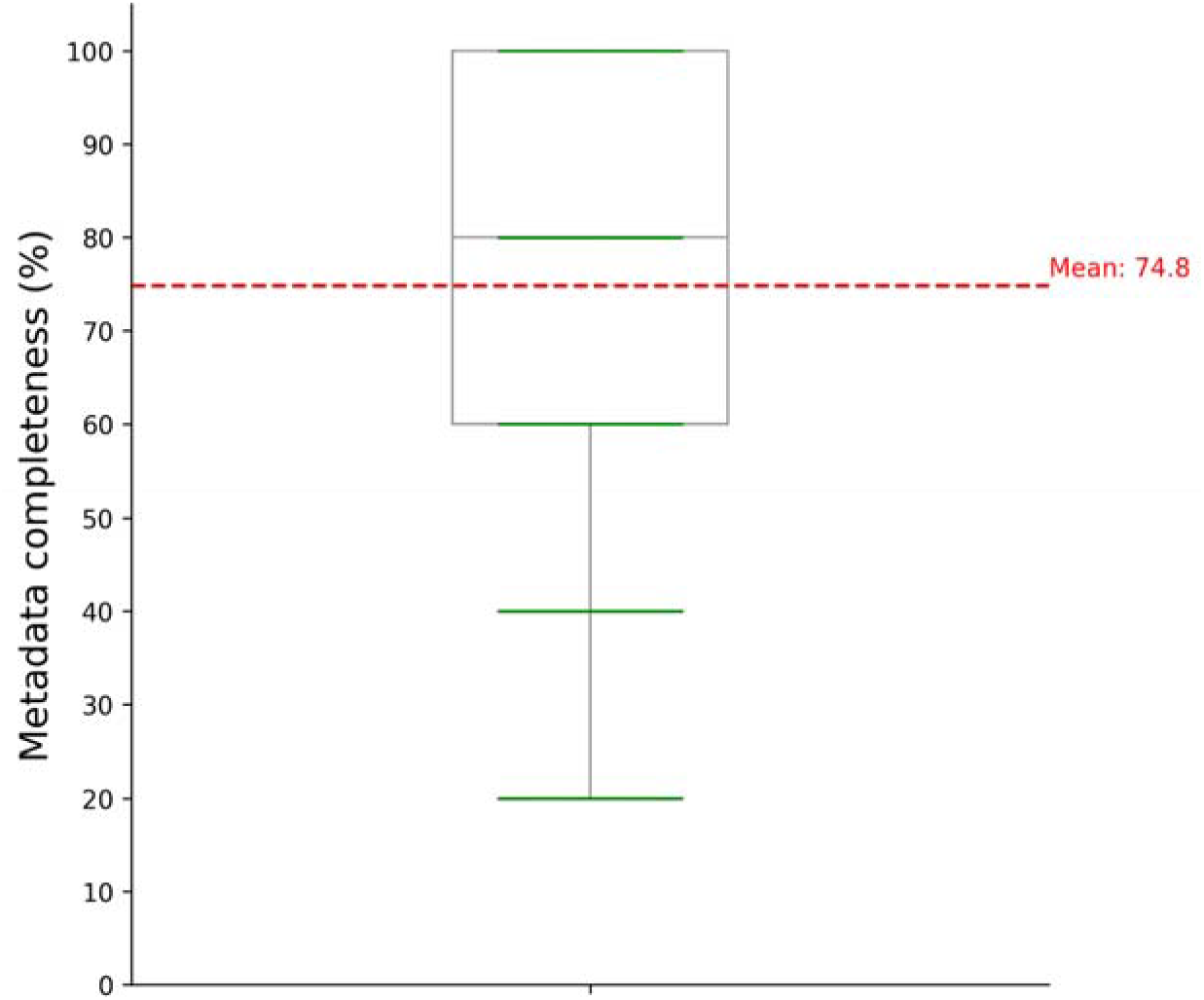
The metadata availability reported in the D1-dataset from the textual content of publications and public repositories. (Red line: the mean metadata availability among the samples in the D1-dataset.)

**Figure S3.**
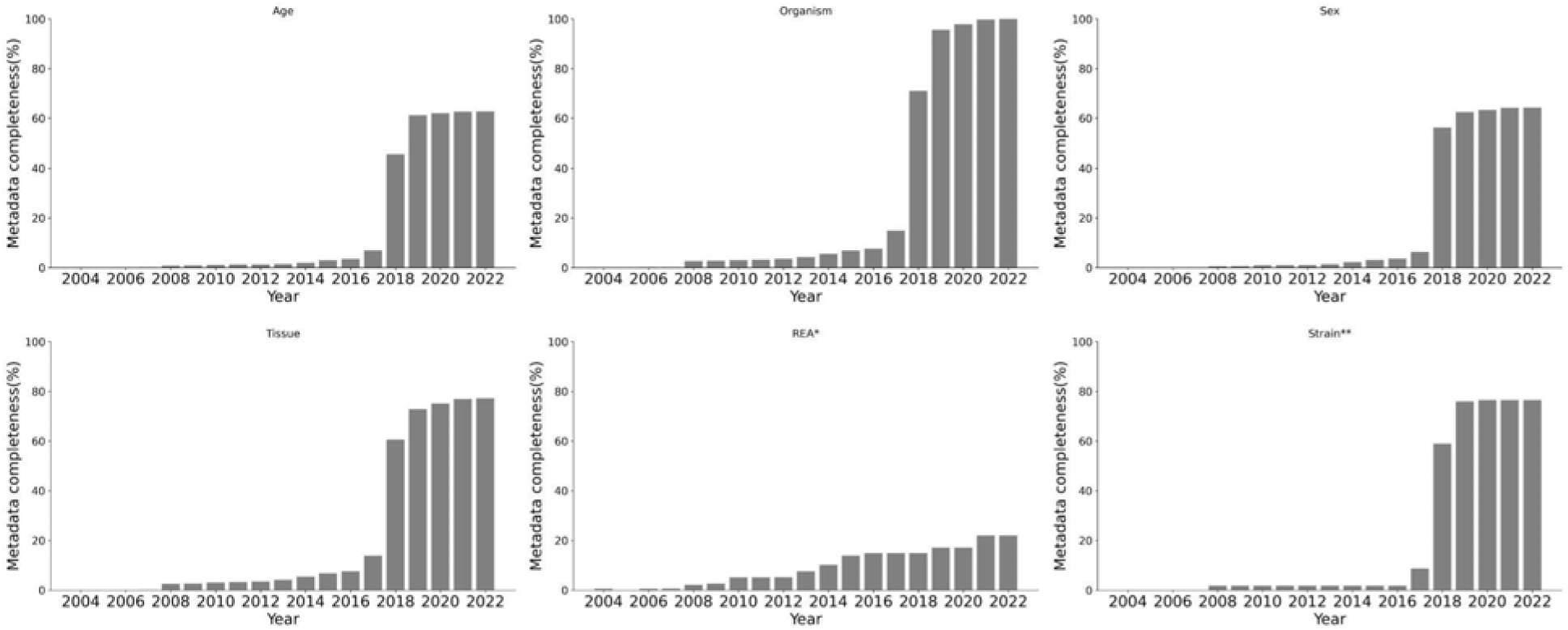
The cumulative metadata availability of the six essential phenotypes in the D1-dataset over the years.

**Figure S4.**
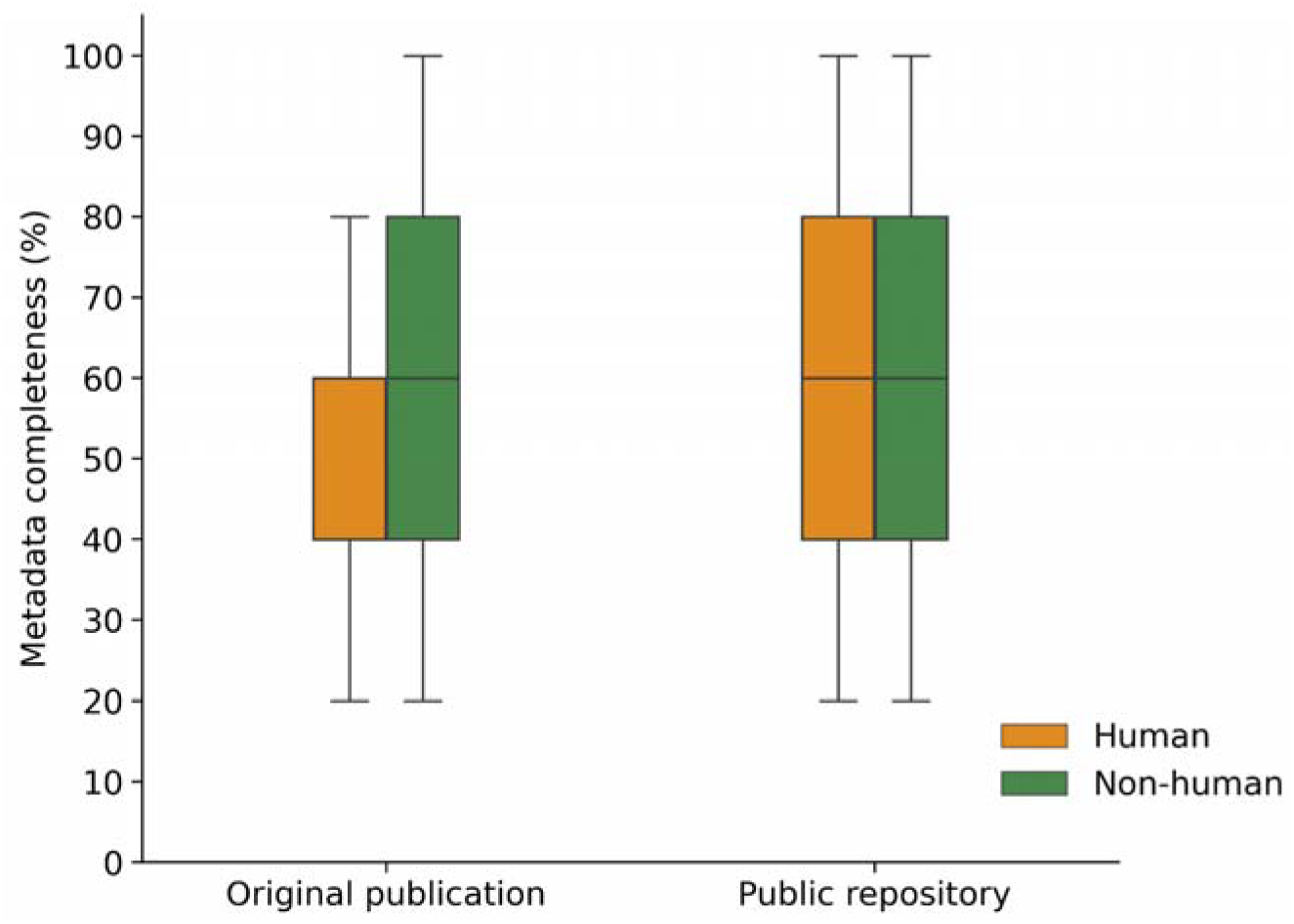
The human and non-human metadata availability in the textual content of publications and public repositories for the D1-dataset.

**Figure S5.**
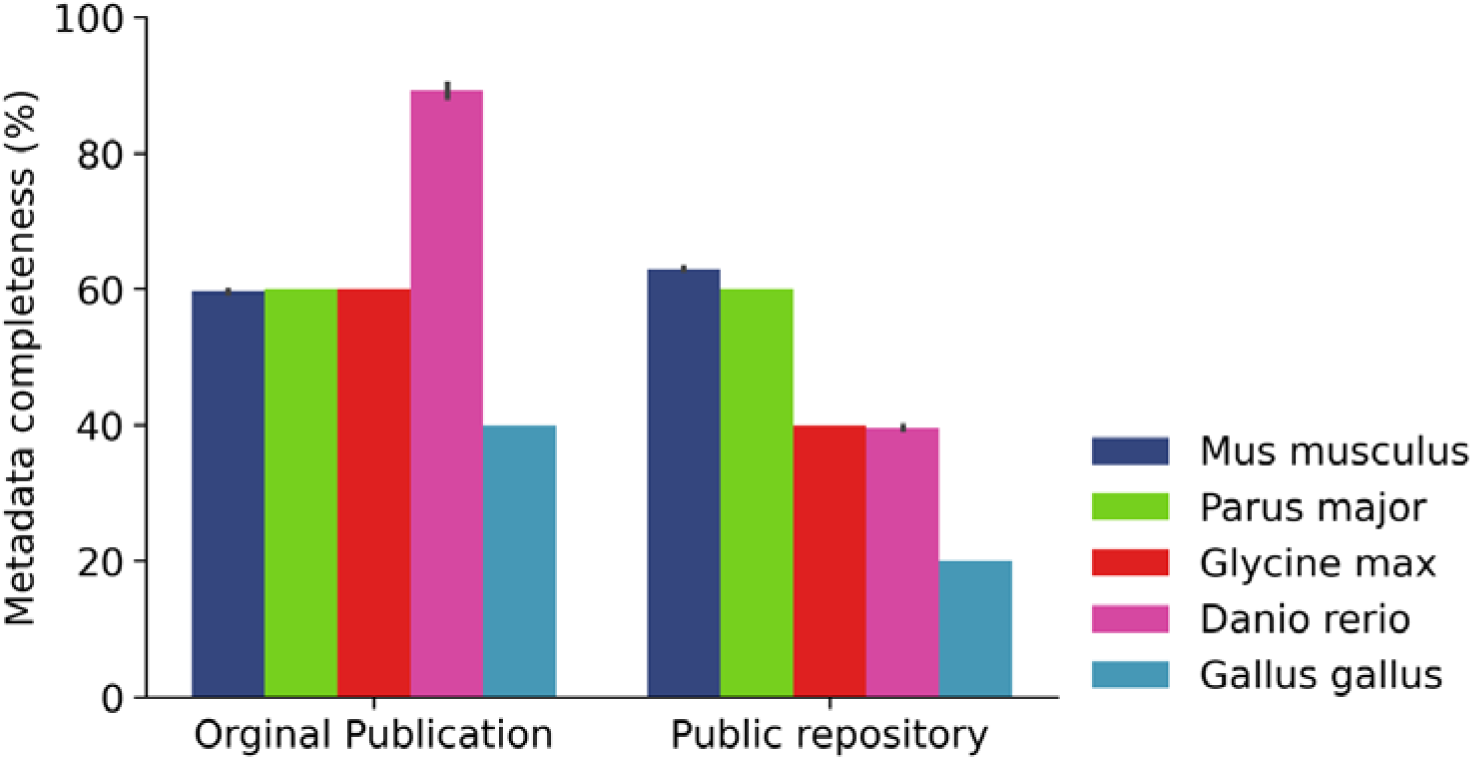
The metadata availability between the textual content of publications and public repositories among the top five non-human organisms represented in the D1 dataset.

**Figure S6.**
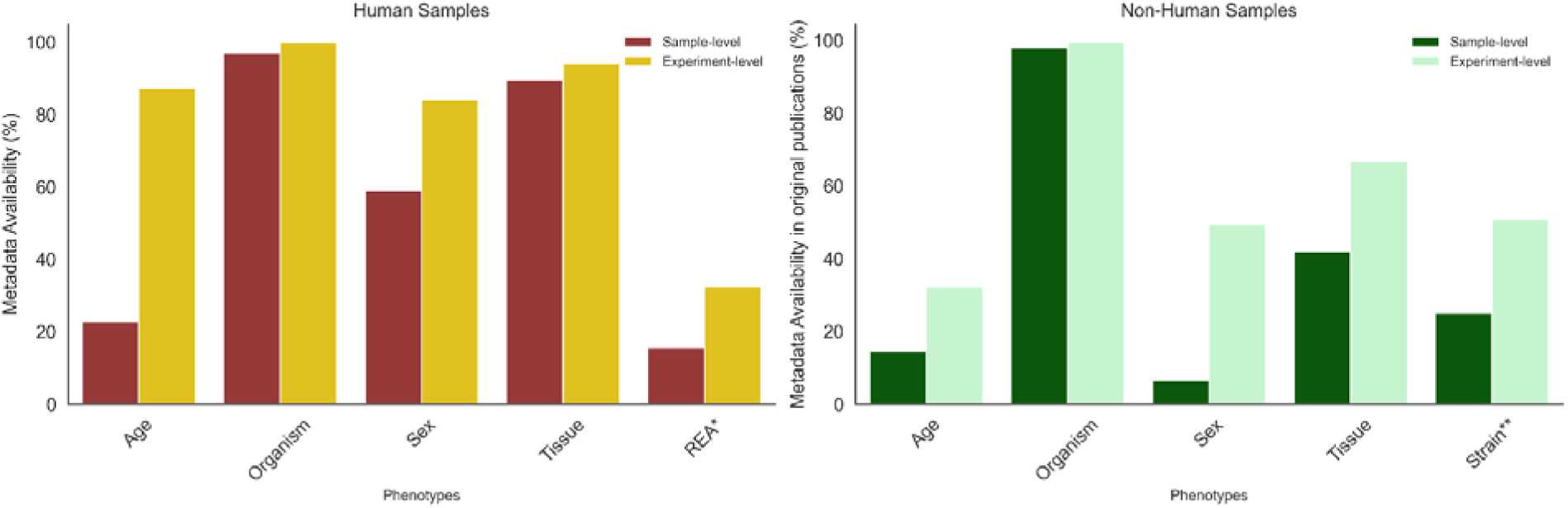
The comparison of the availability of sample-level and experiment-level metadata between human samples (six phenotypes) and non-human samples (five phenotypes) in the D1-dataset.

**Figure S7.**
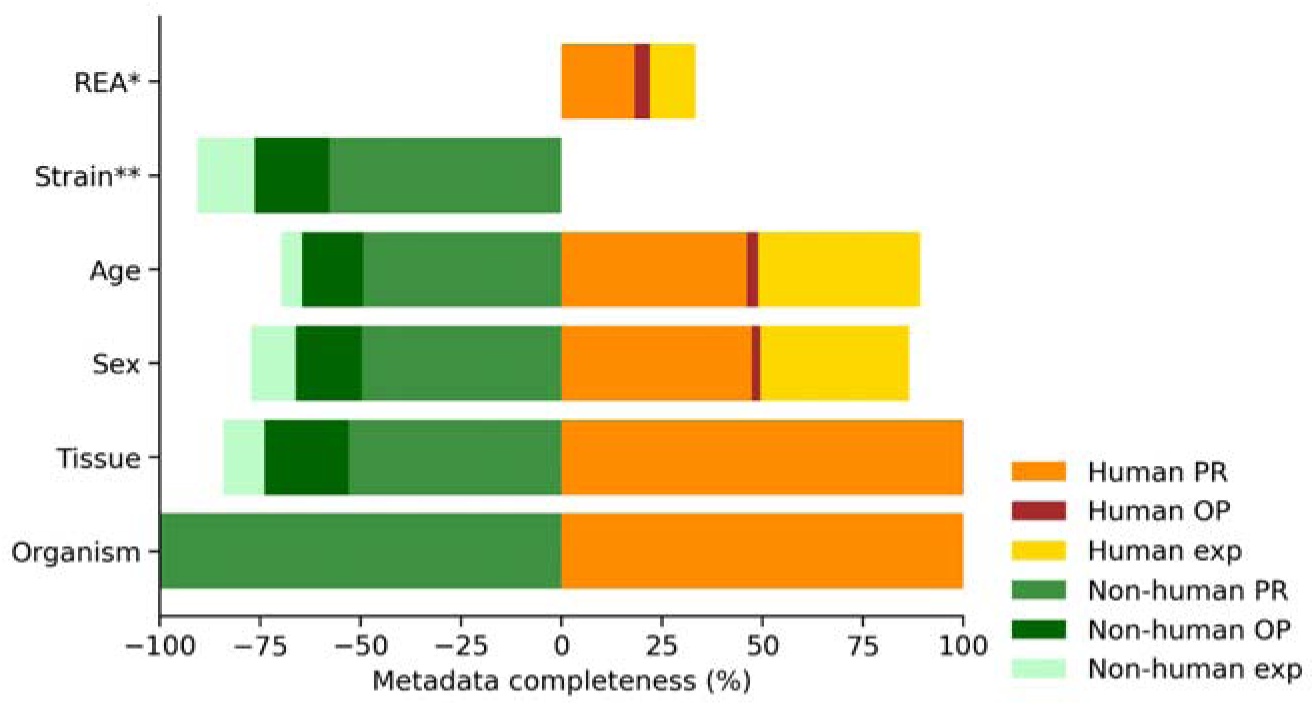
Right: The overall composition of sample-level and experiment-level metadata availability across six phenotypes in the textual content of publications and public repositorie across the human samples in the D1-dataset; Left: The overall composition of sample-level and experiment-level metadata availability across five phenotypes in the textual content of publicationss and public repositories across the non-human samples in the D1-dataset.

**Figure S8.**
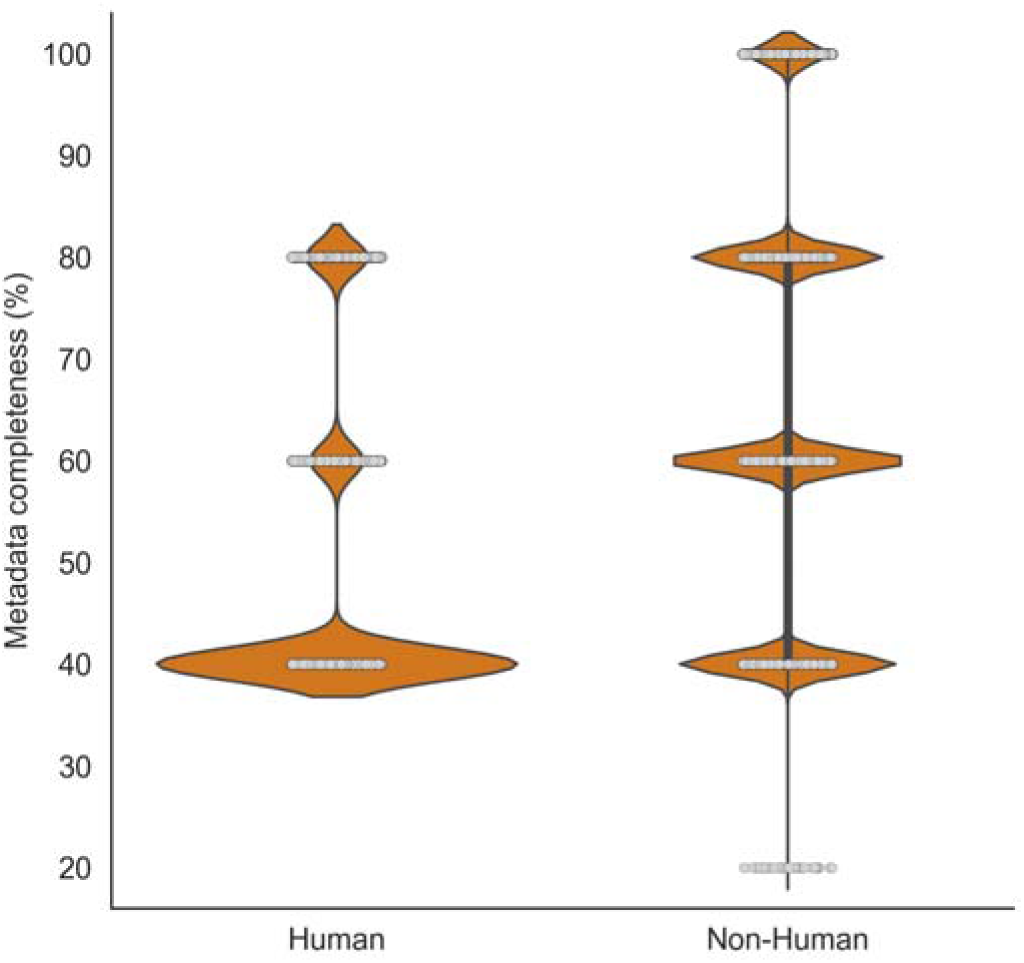
The comparison of the human and non-human metadata availability in the D2-dataset.

**Figure S9.**
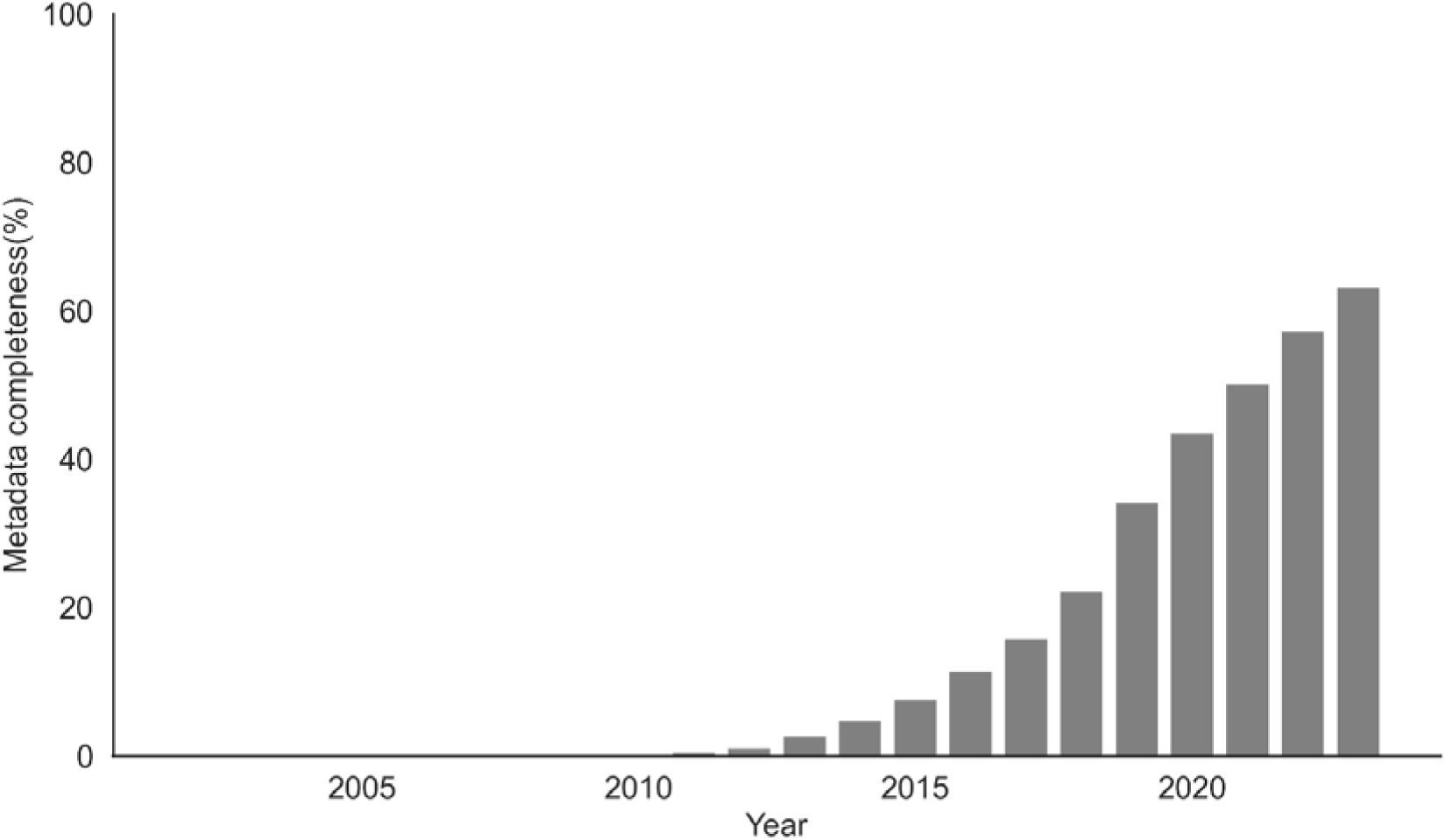
The cumulative metadata availability of the D2-dataset over the years.

**Figure S10.**
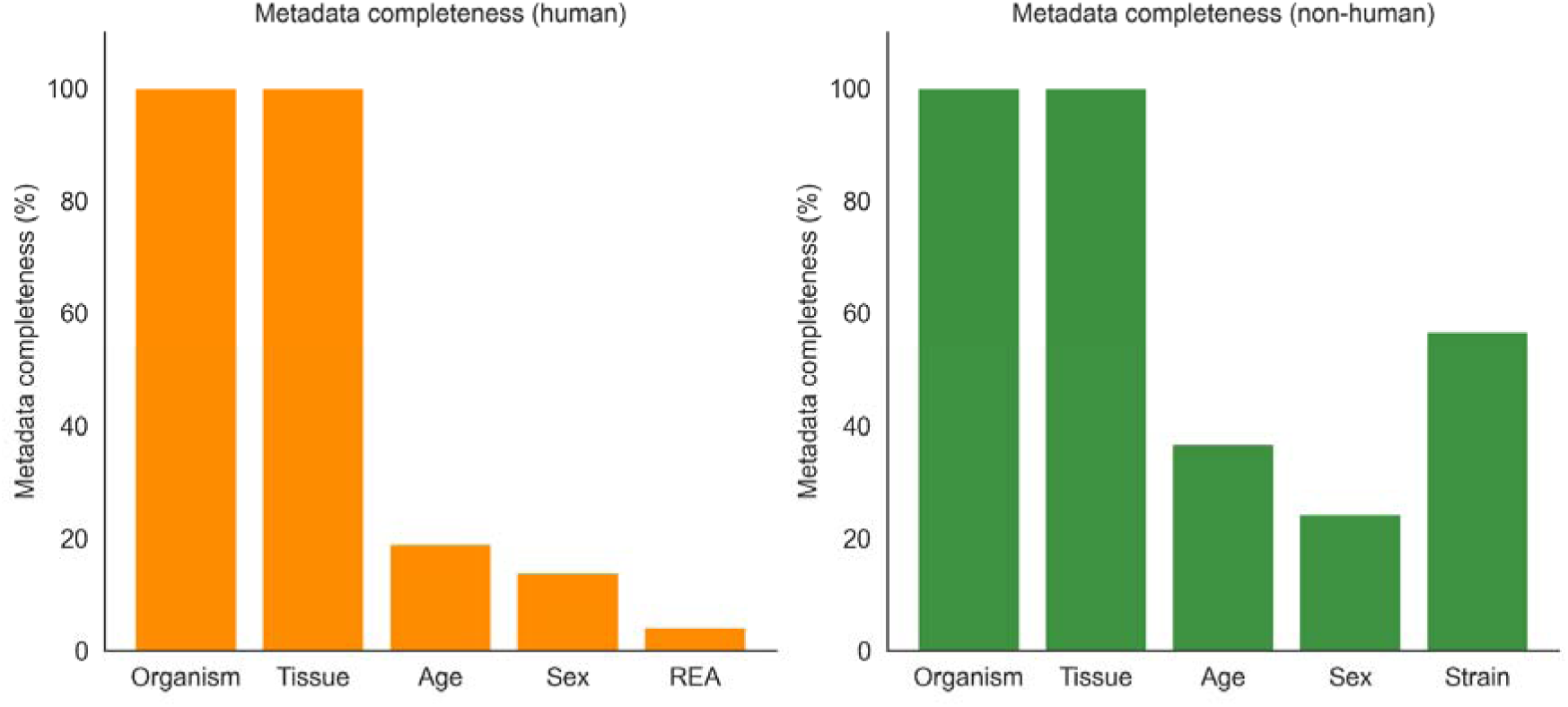
The separate metadata availability of human and non-human samples in the D2-dataset.

**Figure S11.**
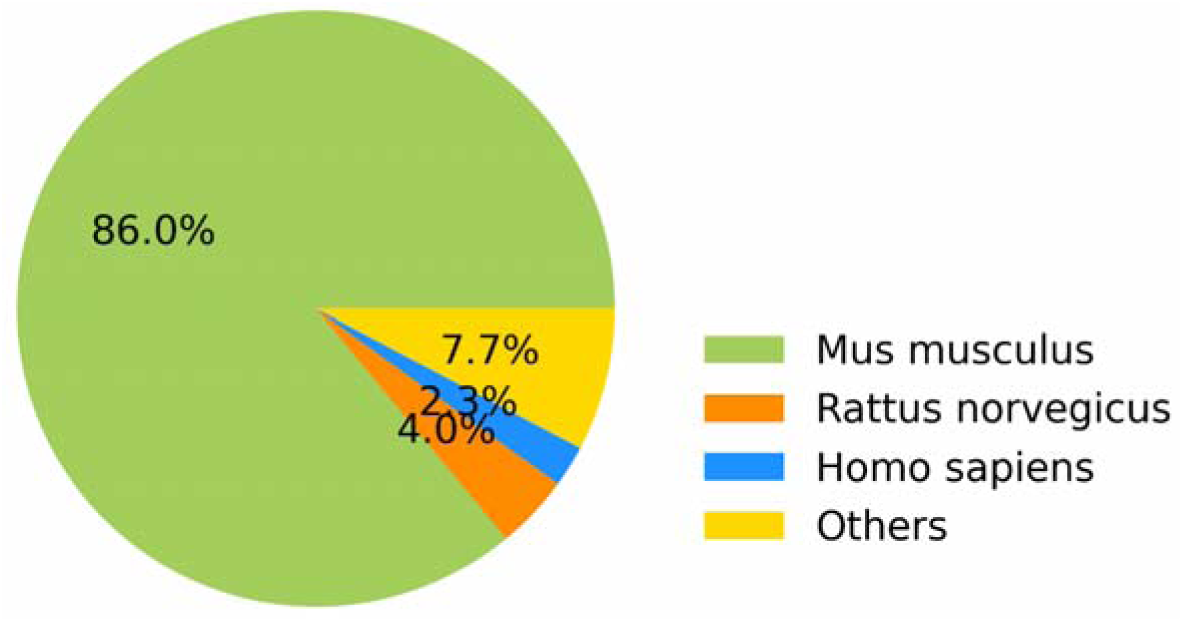
The proportion of organisms among the 2,168,620 samples of the 61,950 studies from the D2 dataset.

