## Supplemental Figures for "The systematic assessment of completeness of public metadata accompanying omics studies in the Gene Expression Omnibus"

**Supplementary Figures**

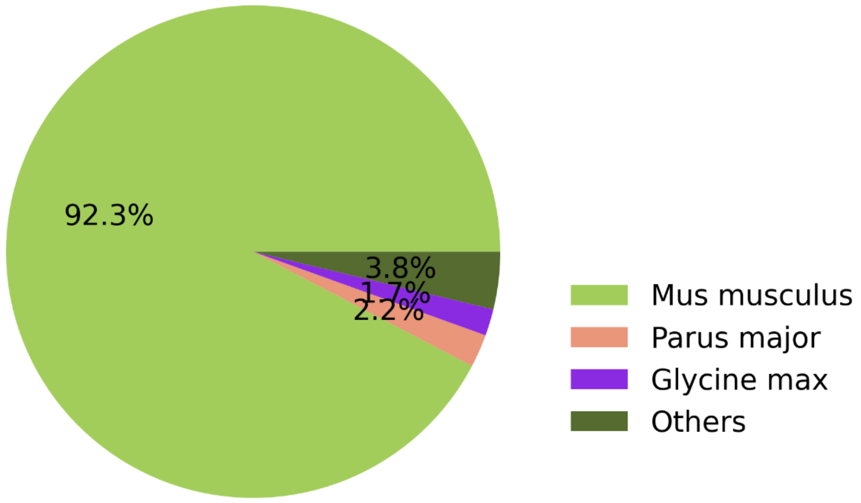


Figure S1. The proportion of non-human samples across 100 studies encompassing 144,862 samples in the D1 dataset.

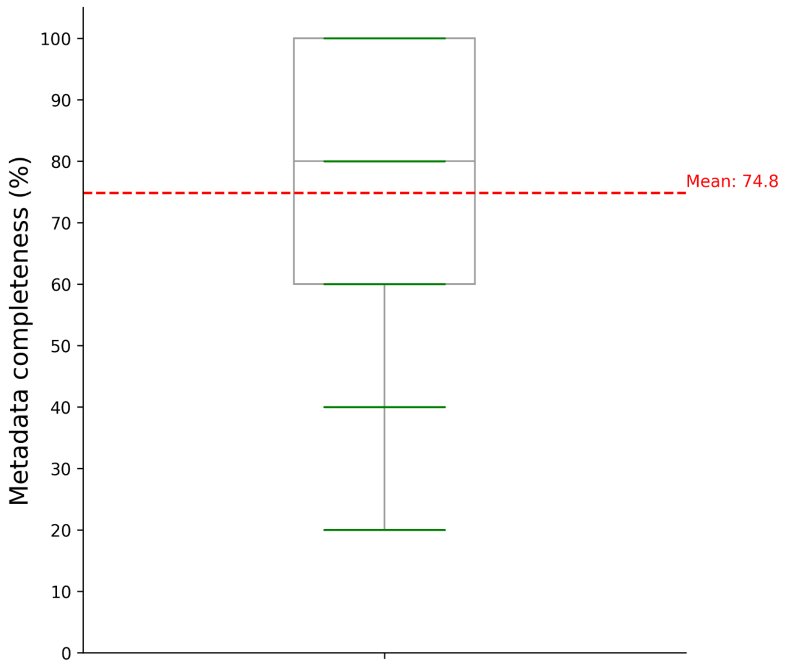


Figure S2. The metadata availability reported in the D1-dataset from the textual content of publications and public repositories. (Red line: the mean metadata availability among the samples in the D1-dataset.)

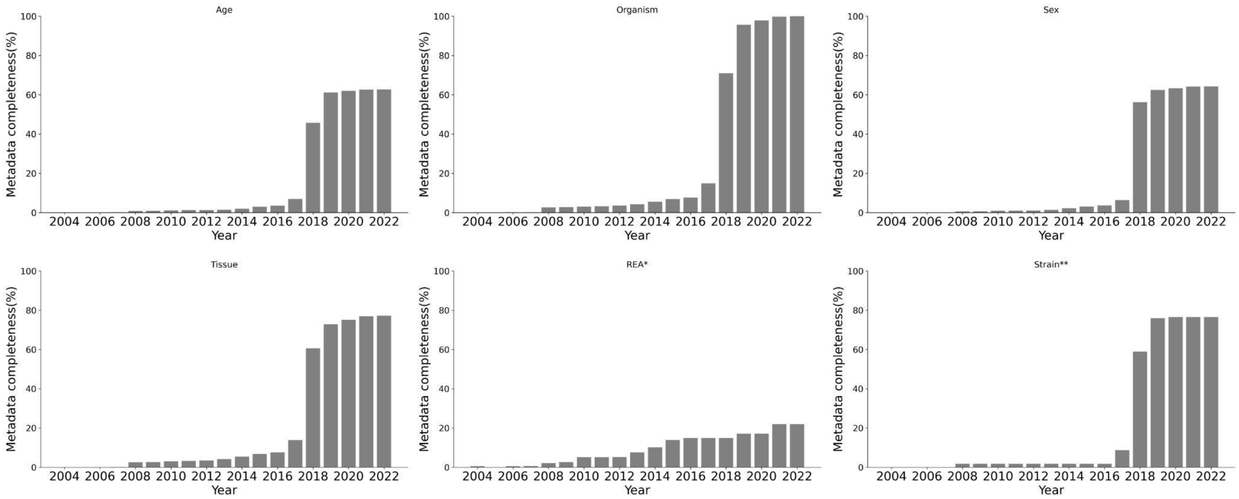


Figure S3. The cumulative metadata availability of the six essential phenotypes in the D1-dataset over the years.

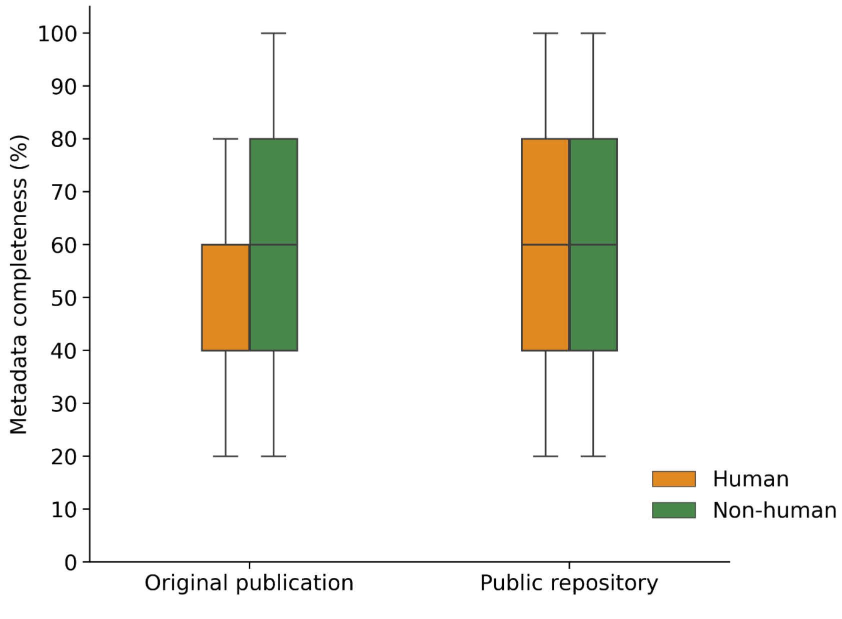

Figure S4. The human and non-human metadata availability in the textual content of publications and public repositories for the D1-dataset.

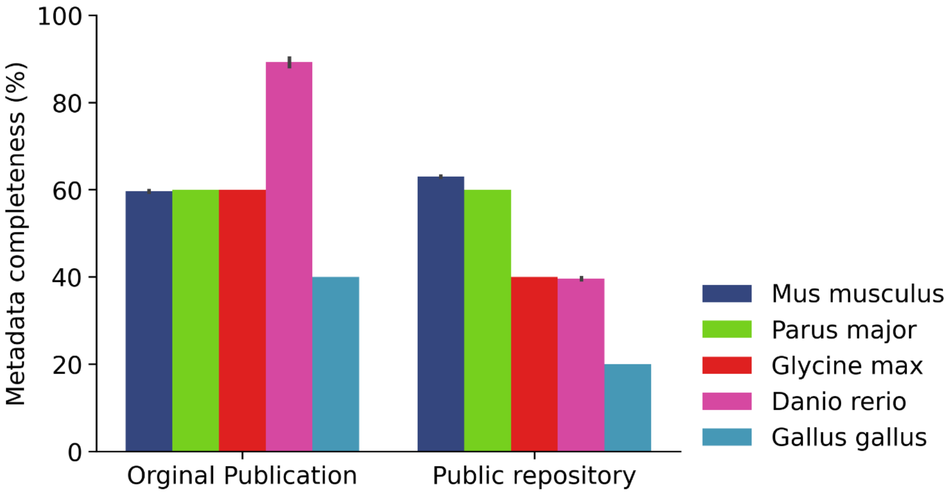


Figure S5. The metadata availability between the textual content of publications and public repositories among the top five non-human organisms represented in the D1 dataset.

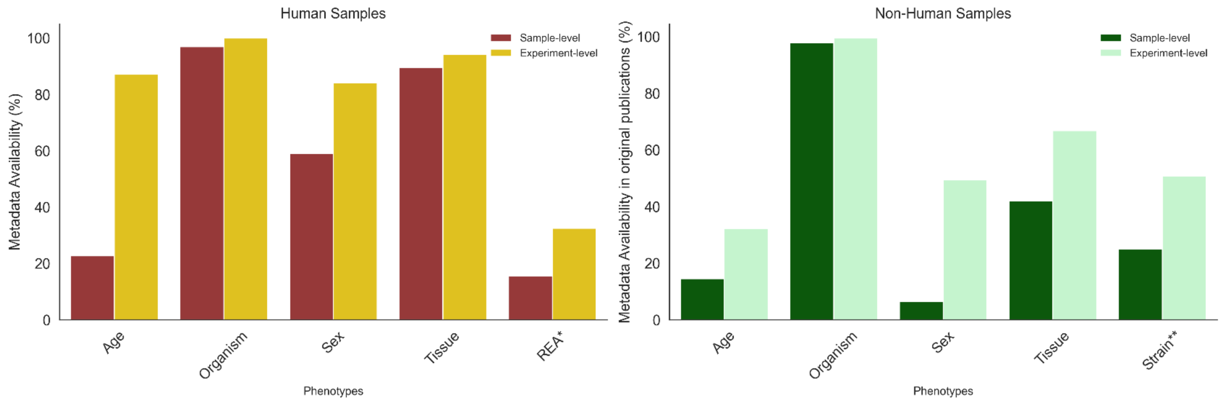


Figure S6. The comparison of the availability of sample-level and experiment-level metadata between human samples (six phenotypes) and non-human samples (five phenotypes) in the D1-dataset.

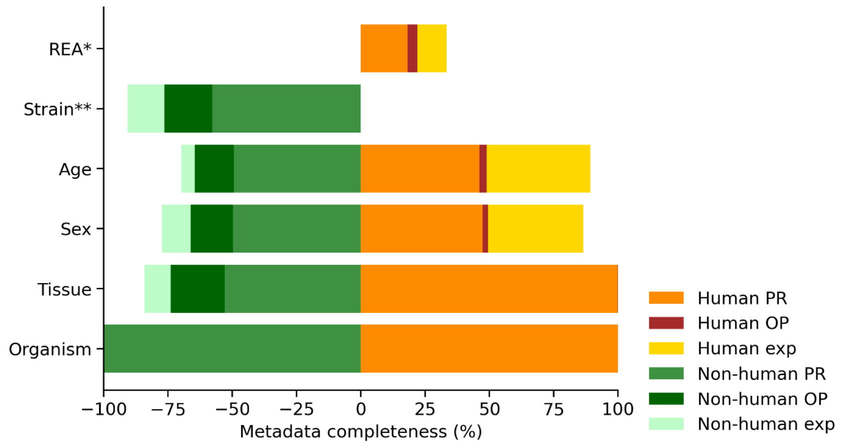


Figure S7. Right: The overall composition of sample-level and experiment-level metadata availability across six phenotypes in the textual content of publications and public repositories across the human samples in the D1-dataset; Left: The overall composition of sample-level and experiment-level metadata availability across five phenotypes in the textual content of publicationss and public repositories across the non-human samples in the D1-dataset.

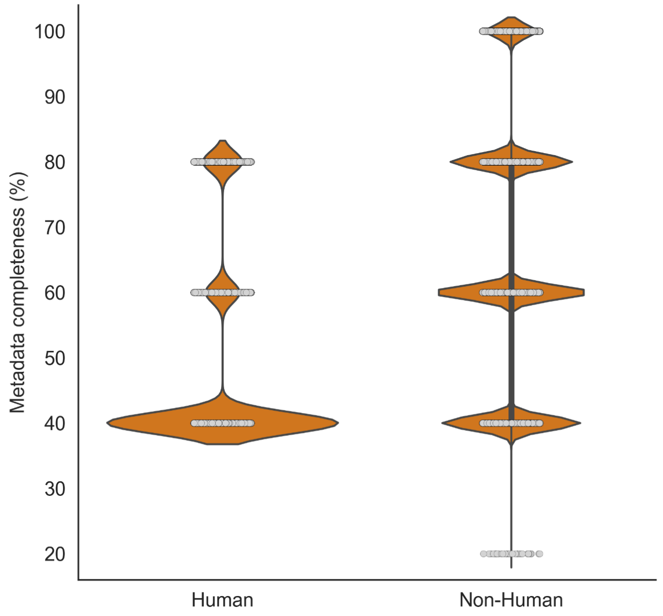


Figure S8. The comparison of the human and non-human metadata availability in the D2-dataset.

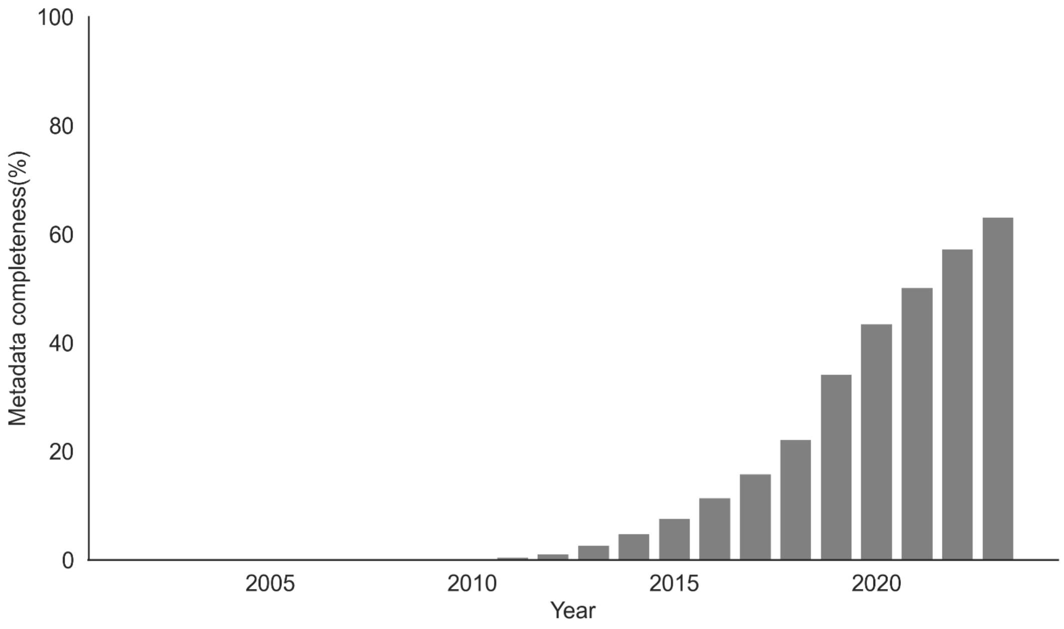


Figure S9. The cumulative metadata availability of the D2-dataset over the years.

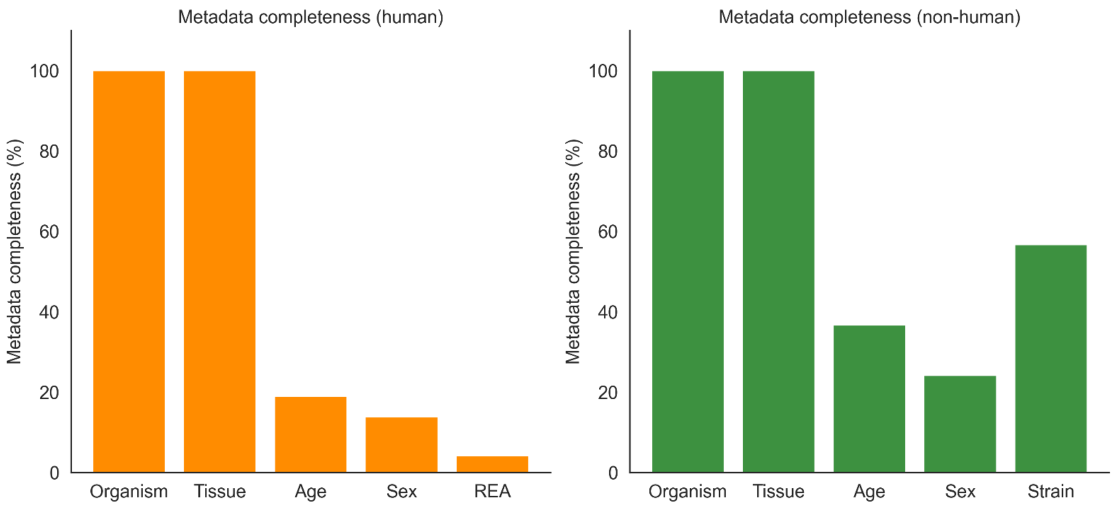


Figure S10. The separate metadata availability of human and non-human samples in the D2-dataset.

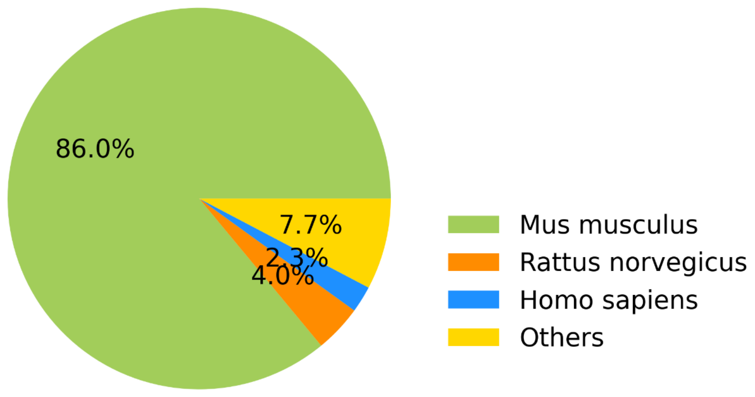


Figure S11. The proportion of organisms among the 2,168,620 samples of the 61,950 studies from the D2 dataset.
